# Large-scale structural variation detection in subterranean clover subtypes using optical mapping validated at nucleotide level

**DOI:** 10.1101/232132

**Authors:** Yuxuan Yuan, Zbyněk Milec, Philipp E. Bayer, Jan Vrána, Jaroslav Doležel, David Edwards, William Erskine, Parwinder Kaur

**Author notes:** Correspondence: Dr. Parwinder Kaur.

## Abstract

Whole genome sequencing has been widely used to detect structural variations (SVs). However, the limited single molecule size makes it difficult to characterize large-scale SVs in a genome because they cannot fully cover such vast and complex regions. Recently, optical mapping in nanochannels has provided novel resolution to detect large-scale SVs by comparing the physical location of the nickase recognition sequence in genomes. Other than in humans, SVs discovered in plants by optical mapping have not been validated. To assess the accuracy of SV calling in plants by optical mapping, we selected two genetically diverse subspecies of the *Trifolium* model species, subterranean clover *cvs*. Daliak and Yarloop. The SVs discovered by BioNano optical mapping (BOM) were validated using Illumina short reads. In the analysis, BOM identified 12 large-scale regions containing deletions and 19 containing insertions in Yarloop. The 12 large-scale regions contained 71 small deletions when validated by Illumina short reads. The results suggest that BOM could detect the total size of deletions and insertions, but it could not precisely report the location and actual quantity of SVs in the genome. Nucleotide-level validation is crucial to confirm and characterize SVs reported by optical mapping. The accuracy of SV detection by BOM is highly dependent on the quality of reference genomes and the density of selected nickases.

## 1 Introduction

Structural variations (SVs) are genomic alterations in sequence size, copy number, orientation or chromosomal location between individuals (Feuk et al., 2006). Usually, they are > 1kbp. SVs are important genetic features that enrich genetic diversity and lead to important phenotypes (Escaramis et al., 2015). Before the advent of molecular biology and DNA sequencing, SVs could only be characterized by cytogenetic analyses (Feuk et al., 2006). The consequent low throughput and low identification rate impeded the understanding of SVs (Saxena et al., 2014). Recently, an increasing number of studies on humans has shown that SVs contribute significantly to the generation of diseases (Feuk et al., 2006;Sharp et al., 2006;Stankiewicz and Lupski, 2010). Although studies of structural variation on plants has been increasing, challenges remain in the accuracy of SV detection. This is mainly due to the lack of high-quality reference genomes for large and complex plants (Saxena et al., 2014). It is difficult to assemble plant genomes using short sequence reads owing to abundant repeats, polyploidy, and numerous pseudogenes in plant chromosomes (Yuan et al., 2017a). Commonly used short sequence reads cannot fully cover large and complex SV regions, leading a difficulty in large-scale SV detection.

Optical mapping in nanochannels, on the other hand, has provided a novel approach in genome assembly and large-scale SV calling (Cao et al., 2014). Contrasting with traditional DNA sequencing, optical mapping uses specific endonucleases to nick DNA strands, followed by fluorescent labelling and image capture to produce long, single molecule maps to reconstruct genome regions (Schwartz et al., 1993). The single molecule maps are >200 kbp on average, which is substantially longer than those DNA single molecules (typically 100 bp to 10 kbp) produced by commonly used DNA sequencing platforms such as Illumina and PacBio platforms (Yuan et al., 2017a).

By mapping the physical locations of nicking sites in reference and query genomes, optical mapping uses query genomes and/or consensus maps (similar to contigs in next generation sequencing, here there are consensus optical maps) to detect SVs by examining the physical location differences, orientation, and multi-alignments between coupled restriction sites. However, it is uncertain whether those detected SVs exist at a nucleotide level or are misreported due to the limited density of nicking sites. To address these concerns, we selected the *Trifolium* model species, subterranean clover (*Trifolium subterraneum* L.), with a high-quality reference and high-resolution BioNano optical maps (BOM) for two genetically diverse subspecies. These BOM findings were then validated by high coverage Illumina short read data generated for the two subtypes.

Subterranean clover is the key forage legume in Australia, producing valued feed for livestock on a sown area of more than 29 million hectares (Nichols et al., 2013). As with other legumes, symbiotic nitrogen fixation in subterranean clover contributes to soil improvement. Subterranean clover is diploid (2n = 2x = 16) with a genome size around 556 Mb/1C. Its inbreeding nature, annual habit, and well-assembled reference genome (*subterraneum*) have established it as a model for *Trifolium* (Nichols et al., 2013;Kaur et al., 2017). Based on morphology, genetic, and cytogenetic data, subterranean clover is classified into three subspecies: *subterraneum*, *yanninicum* and *brachycalycinum* (Katznelson and Morley, 1965a;b). The subspecies differ morphologically, enabling them to adapt to different soil environments, e.g. ssp. *subterraneum* and ssp. *yanninicum* are adapted to moderately acidic soils, with ssp. *subterraneum* found on well-drained soils and ssp. *yanninicum* adapted to water-enriched soils (Francis and Devitt, 1969). In contrast, ssp. *brachycalycinum* is adapted to dry and neutral-to-alkaline soils that contain cracks or stones facilitating burr development. In this study, we examined the sympatric subspecies *subterraneum* and *yanninicum* to check the performance of optical mapping in SV detection and validate the findings using short read sequencing.

## 2 Material and Methods

### 2.1 Purification of cell nuclei

Suspensions of intact cell nuclei were prepared following Vrána *et al.* (Vrana et al., 2016). Approximately 20 g each of mature dry seeds of ssp. *subterraneum* cultivar Daliak and ssp. *yanninicum* cultivar Yarloop were germinated at 25°C on moist paper towels in a dark environment. When the roots reached 2–3 cm in length, they were excised about 1 cm from the root tip, fixed in (2% v/v) formaldehyde at 5°C for 20 min, and subsequently washed three times with Tris buffer (5 min each time). The root tips (~40/sample) were excised and transferred to 1 ml IB buffer (Šimková et al., 2003), in which cell nuclei were isolated using a homogenizer at 13,000 rpm for 18 s. Large debris was removed by filtering through 50-μm nylon mesh, and the nuclei in suspension were stained with DAPI (2 μg ml^−1^).

### 2.2 Preparation of high molecular weight (HMW) DNA

High molecular weight (HMW) DNA was prepared according to Šimková et al. (Šimková et al., 2003) with modifications. Four batches of 700,000 G1-phase nuclei each were sorted into 660μl IB buffer in 1.5 ml polystyrene tubes using a FACSAria II SORP flow cytometer and sorter (BD Biosciences, San Jose, USA). One 20 μL agarose miniplug was prepared from each batch of nuclei. The miniplugs were treated by proteinase K (Roche, Basel, Switzerland), washed in wash buffer (10 mM Tris, 50 mM EDTA, pH 8.0) four times, and subsequently five times in TE buffer (10 mM Tris,1 mM EDTA, pH 8.0). After the plugs had been melted for 5 min at 70°C and solubilized with GELase (Epicentre, Madison, USA) for 45 min, DNA was purified by drop dialysis against TE buffer (Merck Millipore, Billerica, USA) for 90 min.

### 2.3 Construction of BioNano optical map

The latest genome assembly of *T. subterraneum* (*cv*. Daliak) (Kaur et al., 2017) was used as a reference and digested *in silico* using Knickers (v1.5.5). Four available nickases (*Nt.BspQI:* GCTCTTC, *Nb.BbvCI:* CCTCAGC, *Nb.BsmI:* GAATGC, *Nb.BsrDI*: GCAATG) were used to check the frequency of enzyme restriction sites in the reference genome with *Nt.BspQI*, being the most appropriate enzyme to nick the HMW DNA with the expected frequency of 7.1 sites per 100 kbp. In all BioNano experiments, *Nt.BspQI* was used. The DNA was labeled and stained following the manufacturer’s NLRS protocol as described in Kaur et al. (2017). Four runs on the BioNano Irys^®^ instrument (30 cycles/run) were carried for subspecices *yannicum* (*cv*. Yarloop) to achieve sufficient genome coverage (~425X).

The dedicated BioNano IrysView (v2.5.1.29842), BioNano tools (v5122), BioNano scripts (v5134) and runBNG (Yuan et al., 2017b) were used to *de novo* assemble *cv*. Yarloop single molecule optical maps. Before *de novo* assembly, molecule quality was checked by running the ‘Molecule Qlty Report (MQR)’ in BioNano IrysView using *cv*.Yarloop raw BOM data and the digested reference genomes. In the alignment parameter settings, the *p*-value (–T) was set to 1.81 × 10^−08^ and the number of iterations (–M) was set to 5. On receipt of the MQR, we adjusted the *de novo* assembly parameter settings from the default false positive density (–FP) 1.5 to 1.67, default negative rate (–FN) 0.15 to 0.09, default scalingSD (–*sd*) 0.0 to 0.25, default siteSD (–*sf*) 0.2 to 0.15, and default initial assembly *p*-value (–T) 1 × 10^−9^ to 1.81 × 10^−08^.

### 2.4 Structural variation detection by BOM validated using Illumina short reads

After *de novo* assembly, runBNG was used for SV calling. To check the accuracy of the SVs detected by BOM, we selected short paired-end reads for validation. The plants were grown in the field at Shenton Park, Western Australia (31°57’ S, 115°50’ E) and the genomic DNA was extracted from a single plant from each of the two *cv*. Yarloop and *cv*. Daliak of the subterranean clover subtypes. Truseq Illumina libraries were prepared with an insert size of approximately 550 bp and the short paired-end reads were generated using Illumina Hiseq 2000 at coverage of 48× in *cv*. Yarloop and 56× in *cv*. Daliak (the same dataset used in (Kaur et al., 2017)). Reads from both cultivars were aligned to the latest nucleotide reference (*cv.* Daliak) respectively (Kaur et al., 2017) using BWA-MEM (v0.7.12) (Chiang et al., 2015). Results were visualized using the integrative genomics viewer (IGV) (v2.3.91) (Robinson et al., 2011). Nucleotide-level SV calling was performed using Lumpy (v0.2.11) (Layer et al., 2014) and Speedseq (v0.1.0) (Chiang et al., 2015). The settings of Speedseq were the deafult. The program used from Lumpy was ‘lumpyexpress’. The nucleotide reference was the same one used in the BWA-MEM reads mapping. The short sequence reads of *cv*. Yarloop were the same as used in reads mapping. The SV calling was in the whole genome. With large-scale regions containing SVs from BOM identified, we checked the corresponding regions and see if those regions contain SVs from the result of Lumpy and the visualization of IGV.

## 3 Results

### 3.1 *De novo* assembly of cv. Yarloop optical map

A total of 1,083,671 single molecule maps (raw optical maps) was generated from the BioNano Irys platform with a total length of 235.5 GB (~425× genome coverage), of which the molecule N50 was 212.7 kbp, and the average label density was 7.5 per 100 kbp (Table 1). After filtering out low-quality single molecule maps using the default setting (<150 kbp), 958,136 single molecule maps remained with a total length of 212.7 GB (~385× genome coverage), of which the molecule N50 was 218.6 kbp, and the average label density was 8.3 per 100 kbp. Using the filtered single molecule maps, 375,975 single molecule maps were *de novo* assembled to each other to generate 377 consensus maps. The total length of the generated consensus maps was 475 Mb (~89% of the total length of the reference genome) with a map N50 of 1.8 Mb.

**Table 1.** Statistics of *cv*. Yarloop BioNano optical maps

| Subject | Raw BioNano data | Filtered BioNano data | Assembled BioNano data |
| --- | --- | --- | --- |
| Number of molecules | 1,083,671 | 958,136 | 375,975 |
| Number of consensus maps | N/A | N/A | 377 |
| Total length | 235.5 Gb | 212.7 Gb | 475.2 Mb |
| N50<sup>□</sup> | 212.7 kbp | 218.6 kbp | 1.8 Mb |
| Average of label density (/100 kbp) | 7.5 | 8.3 | N/A |
| Coverage | 425 | 385 | 0.89 |

<sup>□</sup>In the set of molecules, N50 represents the length of the shortest molecule whose length is greater than half of the total sum of lengths of all molecules; it is the point of half of the mass of the distribution

### 3.2 SVs assessment with BOM validated by Illumina short reads

In the BioNano SV calling between *cv*. Yarloop BioNano molecule maps and the *cv.* Daliak reference genome, 12 large-scale regions (tens of kbp regions) containing deletions and 19 containing insertions were identified in *cv*. Yarloop (Supplymentary Figure 1). The average length of the deletions in the 12 regions was estimated as 6.2 kbp (Supplymentary Table S1) and, in these regions, 9.7% of the sequences were assembly gaps (N’s). Regarding insertions, the average length of the insertions was 8.04 kbp in the 19 regions, and the total percentage of unknown sequences in these insertion regions was 3.6%. The Lumpy SV calling detected 20,887 deletions, 115 inversions, and 1,331 duplications in *cv*. Yarloop compared with the 71 detected deletions that supported the 12 regions implied by BOM in *cv*. Daliak (Supplymentary Figure 2 and Supplymentary Table S2). Lumpy did not detect any insertions.

## 4 Discussion

BioNano single molecule maps are substantially longer than those produced by traditional sequencing methods, which means that BioNano single molecule maps can easily cover most of the large and complex genome regions that next generation sequence reads cannot span. In the *de novo* assembly of *cv*. Yarloop BioNano single molecule maps, the total length of the consensus maps accounted for ~89% of the estimated *T. subterraneum* genome size contrary to our expectation of ~100%. This incomplete assembly could be caused by the low-quality single molecule map filtering step or the single map fragmentation due to the close proximity of *Nt.BspQI* restriction sites leading to DNA double-strand breaks in some DNA regions (Hastie et al., 2013). These fragmented single molecule maps may collapse during *de novo* assembly causing assembly problems. Alternatively, some repetitive regions in *cv*. Yarloop might be longer than the length of BioNano single molecule maps which may have collapsed during *de novo* assembly.

By aligning consensus genome maps to a reference, BioNano Genomics uses a multiple local alignment algorithm to perform SV calling. SVs are detected as alignment outliers, which are defined by two well-aligned regions flanking poorly aligned or unaligned regions. To avoid false positive in SV calling, BioNano Genomics claims that the algorithm implemented in ‘runSV’ considering the non-normalised p-values of two well-aligned regions and the non-normalised log-likelihood ratio of the poorly aligned or unaligned regions. In the performance of SV calling reported by BioNano Genomics, when the effective coverage for a haplotype-sensitive assembly ≥70 X, the sensitive for homozygous insertions and deletions (≥ 1kbp) is over 98%. In this research, all SVs detected were larger than 1 kbp.

Lumpy is one of the most popular and reliable SV callers using short read sequencing to detect SVs. Different from other SV callers using one signal such as read-pair, split-read, read-depth and prior knowledge, to detect SVs, Lumpy integrates multiple SV signals and uses a probabilistic framework to increase the sensitivity in SV calling (Layer et al., 2014). In the Lumpy SV calling, we identified 71 small deletions in the 12 large-scale regions reported by BOM. While, the total length of the 71 deleted genomic regions reported by Lumpy was close to the total length reported by BOM (71.7 kbp *vs.* 74.4 kbp respectively), some length differences remained, probably due to the incorrect gap size or misassemblies in the reference genome, or also could be due to the incomplete SV calling in Lumpy. Interestingly, the gaps in the detected SV regions which were highly likely caused by collapse in the repetitive regions, was complemented by BioNano super-scaffolding process for the generation of the advanced reference assembly (Kaur et al., 2017).

No insertions were reported by Lumpy in the SV calling, probably those sequences being novel in *cv*. Yarloop compared to the reference assembly based on the *cv*. Daliak. When nucleotide level alignments were carried out using short sequence reads from the *cv*. Yarloop with the *cv*. Daliak, Yarloop reads from genomic regions not present in the reference assembly, either being Yarloop-specific or unassembled in the reference could not map. As such, SV could not be called in these regions. Those novel sequences were grouped as unmapped sequences, earlier abandoned by Lumpy in SV calling. This issue has also been reported previously by Xia *et al*. (Xia et al., 2016) for most reference based SV calling methods, which cannot efficiently report large-scale insertions if there are many novel sequences in the examined individuals.

In terms of SV calling, in this study BOM identified fewer SVs than those reported by Lumpy using whole genome sequencing (Figure 1). The location of SVs detected by BOM is only approximate. The precise location of SVs inside the reported regions is uncertain in the absence of other long range sequencing data. Although BOM can report the size of SVs, owing to the density of enzyme restriction sites in the range of 10 kbp, the size is more likely a size aggregation of several small deletions (see SV size comparison between SVs called by BOM and Illumina short reads in Supplementary Figure 2. Small deletions are those DNA regions with a length from few base pair to few hundred base pair). Incorrectly placed/oriented contigs/scaffolds and incorrect estimates of gap sizes between contigs can also affect SV detection in optical mapping, particularly for deletion and insertion, as it is based on the recognition site of the nickase(s) used.. Inaccurate gap size has a high probability to call false positive SVs. Gap regions may contain enzyme restriction sites that cannot be represented in the reference genome, leading to mismatches or missing reports of SVs. Furthermore, misassemblies can confound the alignment of enzyme restriction sites between maps and report false positive SVs. Clearly, a high-quality reference genome is crucial in the discovery of SVs in BOM.

**Figure 1:**
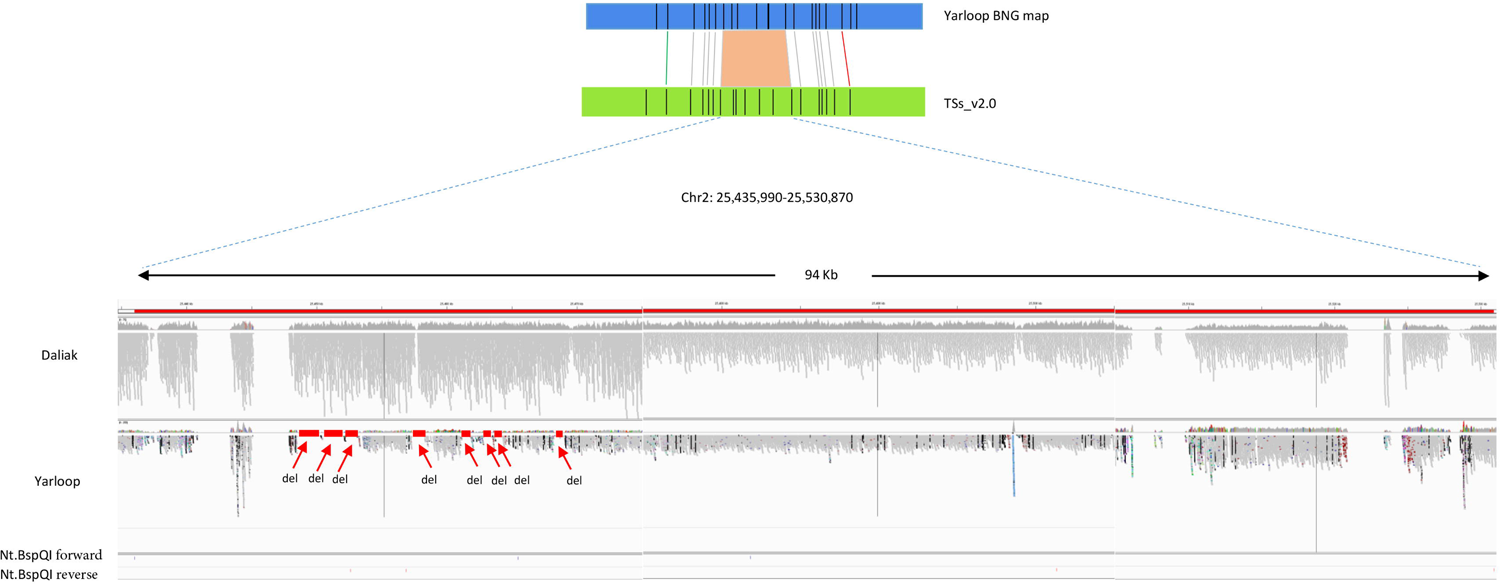
An example of deletions detected by BioNano optical mapping with nucleotide sequences validation. The region reported by BioNano contains deletion(s) in cv. Yarloop compared to the reference genome between location 25,435,990 bp and 25,530,870 bp in chromosome 2. The grey bars in this figure reprsenst short reads aligned to the reference. Other color dots means different SNPs. Nt.BspQI forward reprsents sequence: GCTCTTC and Nt.BspQI reverse repesents sequence: GAAGAGC. The size of the deletion reported by BioNano is 6.6 kbp. From the nucleotide-level validation, eight small deletions (displayed as ‘del’) were visualized in the IGV with no sequence reads aligned to the reference genome. The total size of those eight small deletions is 6.1 kbp. The eight small regions were supported in the Lumpy SV calling.

## 5 Conclusions

Based on the physical location of nicking sites, optical mapping provides an attractive method to detect SVs. Single molecule maps produced by optical mapping are long enough to span most of the large and complex genome regions that traditional sequencing technologies are unable to achieve. However, optical mapping has some limitations in discovering the precise location and actual number of SVs owing to enzyme physical locations. NGS is useful to characterise SVs identified by optical mapping.

Although optical mapping provides the total size of SVs in a detected region, the total size of those SVs can be misreported due to the inaccurate gap size in the reference genome and/or absent enzyme restriction site information in the gap regions. To improve SV detection and characterization, a high-quality reference genome is crucial. In the absence of a high-quality reference genome, possible nucleotide-level validation of those identified SV regions is recommended to assess the accuracy of SV calling in optical mapping.

## 6 Abbreviations

BOM: BioNano optical mapping
DNA: deoxyribonucleic acid
MQR: molecule quality report
N/A: not available
NGS: next generation sequencing
SNP: single-nucleotide polymorphism
SV: structural variation

## 7 Acknowledgments

YY is supported by the China Scholarship Council for his PhD studies at the University of Western Australia. We thank Zdeňka Dubská for assistance with nuclei flow sorting, Helena Staňková for help with BioNano mapping, and Hana Šimková for advice on BioNano mapping. We acknowledge the supercomputing resources provided by the Pawsey Supercomputing Centre with funding from the Australian Government and the Government of Western Australia.

## 8 Author Contributions

KP, ED, BP and YY conceived and designed the research. MZ, VJ and DJ performed the BioNano Irys^®^ System genome mapping experiments. YY performed the bioinformatics analysis, prepared the figures and wrote the manuscript with contributions from KP, BP, MZ, EW, ED, DJ and VJ. All authors read and approved this manuscript.

## 9 Conflict of interest

The authors declare that they have no competing interests.

## 10 Funding

This study was conducted by the Centre for Plant Genetics and Breeding (PGB) at The University of Western Australia (UWA) in close collaboration with Institute of Experimental Botany, Centre of the Region Haná for Biotechnological and Agricultural Research, Czech Republic. This project was also supported by grant award L01204 from the National Program of Sustainability I and by the Czech Science Foundation (award no. P501/12/G090).

## 11 Availability of Data

All raw nucleotide data and BioNano data are under BioProject PRJNA404013.

## 12 Supplementary Material

The supplementary Material for this article can be found in the Supplementary Material for Frontiers.

